# Maintaining performance under pain is effortful: experimental and computational evidence

**DOI:** 10.64898/2026.02.13.705857

**Authors:** Thomas Mangin, Ilaria Monti, Mélysiane Marcotte, Stephane Baudry, Mathieu Roy, Pierre Rainville, Benjamin Pageaux

## Abstract

Pain creates a competing demand on attention, though its impact on performance remains debated. Motivational intensity theory predicts that resource reallocation can preserve output until motivational or capacity limits are reached. Forty adults took part in two preregistered experiments. They completed parallel cognitive (choice reaction, n=20) and motor (isometric hand grip, n=20) tasks at three difficulty levels while receiving warm, low, or high pain heat stimulation on the opposite forearm. Across both domains, participants maintained performance despite painful stimulation. This preservation, however, relied on increased effort mobilization: perceived effort rose with pain intensity, while pain perception decreased during motor and cognitive task execution. Trial wise computational modelling revealed that perceived effort was better predicted by subjective pain experience than by stimulus temperature, supporting a compensatory regulatory mechanism. Thus, maintained performance under pain reflects active resource allocation via effort, generalizable across cognitive and motor domains, but achieved at the cost of increased perceived effort.

## INTRODUCTION

Nociceptive stimulation elicits pain, a salient signal that captures attention and interferes with executive and motor functions (Legrain et al., 2009; Torta et al., 2017). From this perspective, pain is assumed to impair cognitive and motor performance by diverting resources away from task execution. Performance decrements in the presence of pain have been reported across cognitive and motor domains (Bank et al., 2013; Buhle & Wager, 2010; Tabry et al., 2020; Torta et al., 2017); however, these effects are not systematic (Pud & Sapir, 2006; Veldhuijzen et al., 2006). This dissociation suggests that compensatory mechanisms may allow performance to be maintained under certain conditions. However, a key question concerns the extent to which this reallocation of attentional resources is effortful. It may occur with no subjective cost, such that focusing on the task is sufficient and pain modulation emerges automatically as a consequence. Alternatively, this reallocation of resources is active and may entail costs, reflecting a deliberate attempt to ignore pain in order to maintain task focus, and reflected in higher perceived effort (Mangin & Pageaux, 2026). Critically, if pain is actively inhibited, this compensatory process should also reduce subjective pain, yielding a task-induced hypoalgesic effect (Vogel et al., 2022). Distinguishing between these possibilities is critical, as they have different implications for the use of distraction to reduce pain, which may be either universally beneficial or associated with costs. In this view, effort (i.e., intentional engagement of resources) is proposed to be the key variable mediating the relationship between pain and performance.

The motivational intensity theory provides a principled framework to formalize this trade-off (Brehm & Self, 1989; Richter et al., 2016). According to this account, effort reflects the mobilization of resources required to meet task demands and can be flexibly adjusted in the presence of competing demands. For submaximal tasks, increased difficulty, such as induced by pain, can be accommodated through greater effort investment (Richter et al., 2016; Silvestrini & Gendolla, 2019), thereby preserving performance. In contrast, when maximal motivation or capacity is reached, further increases in demand necessarily result in performance decrements or disengagement (Richter et al., 2016; Silvestrini & Gendolla, 2019).

Despite its central role, the determinants of perceived effort remain unclear. Because nociceptive stimulation introduces an additional demand on task execution, it remains unclear whether perceived effort is influenced by objective nociceptive input or subjective pain experience targeted by regulatory processes. Here, perceived effort is considered an index of resource allocation. Disentangling these two accounts requires a formal comparison of their predictions, enabled by computational modeling. To address these questions, we conducted two experiments: one cognitive and one motor, in which task demand (control, low, high) and thermal stimulation (warm, light pain, high pain) were independently manipulated. This design allowed us to test the preregistered hypothesis that performance can be maintained in the presence of pain through compensatory resource allocation. Specifically, we predicted that increasing pain would not systematically impair performance during submaximal task demands but would be associated with a graded increase in perceived effort. We expect that this compensatory process would be accompanied by task-induced hypoalgesia, reflected in reduced subjective pain during task execution relative to control conditions. Finally, to further test our hypotheses, we examined with computational modeling whether perceived effort is better predicted by subjective pain experience than by objective nociceptive stimulation.

## MATERIALS AND METHODS

Forty healthy adults (mean ± SD; age: 22 ± 1.7 years; 20 females) volunteered in the preregistered cognitive (n=20) and motor (n=20) experiments. All procedures were approved by the Ethics Committee of the Montreal Geriatrics Institute.

The preregistrations for both experiments can be found respectively at the following links: [cognitive experiment: osf.io/wf8es; motor experiment: osf.io/5hgwk]. Sensitivity analysis and full details of the methods are in supplementary materials.

### Protocol overview

Both the motor and cognitive study were composed of two laboratory visits spaced by 24h to one week. Both visits took place at the same time of the day (± 1h) to control the influence of the circadian cycle.

During visit 1 (∼2h), participants were asked general questions to control for confounding factors (e.g., substances use such as cigarettes, caffeine, alcohol and drugs, physical activities practiced and the amount of sleep the night preceding the experiment) and they filled out the Pain Catastrophizing Scale Questionnaire. Successively, they went through a sensory calibration of thermal stimulations, a maximal voluntary contraction peak force calculation (only for the motor experiment), a ramp-up of effort with familiarization of the visual analog scale related to the perceptions of effort, and a familiarization with the motor or cognitive task consisting of only 6 trials.

In visit 2 (∼1.5h) participants performed the motor or cognitive task in the presence of thermal stimulation. At the beginning of the visit, temperatures used for the thermal stimulation were validated and adjusted if necessary. Participants then completed either the cognitive or motor task that consisted of 36 15s trials at three levels of difficulty: no demand (control; looking at a fixation cross, or holding the dynamometer), low or high demand (performing a modified Simon task or matching a target at 5% or 30% of their maximal voluntary contraction; see Fig. 1.A). Simultaneously to the execution of each trial, participants received thermal stimulation at one of three individually calibrated temperatures to produce sensations of warmth (control), light or high pain. After each trial, participants reported their perception of the effort required to complete the task, followed by their perception of warmth or pain.

**Figure 1.**
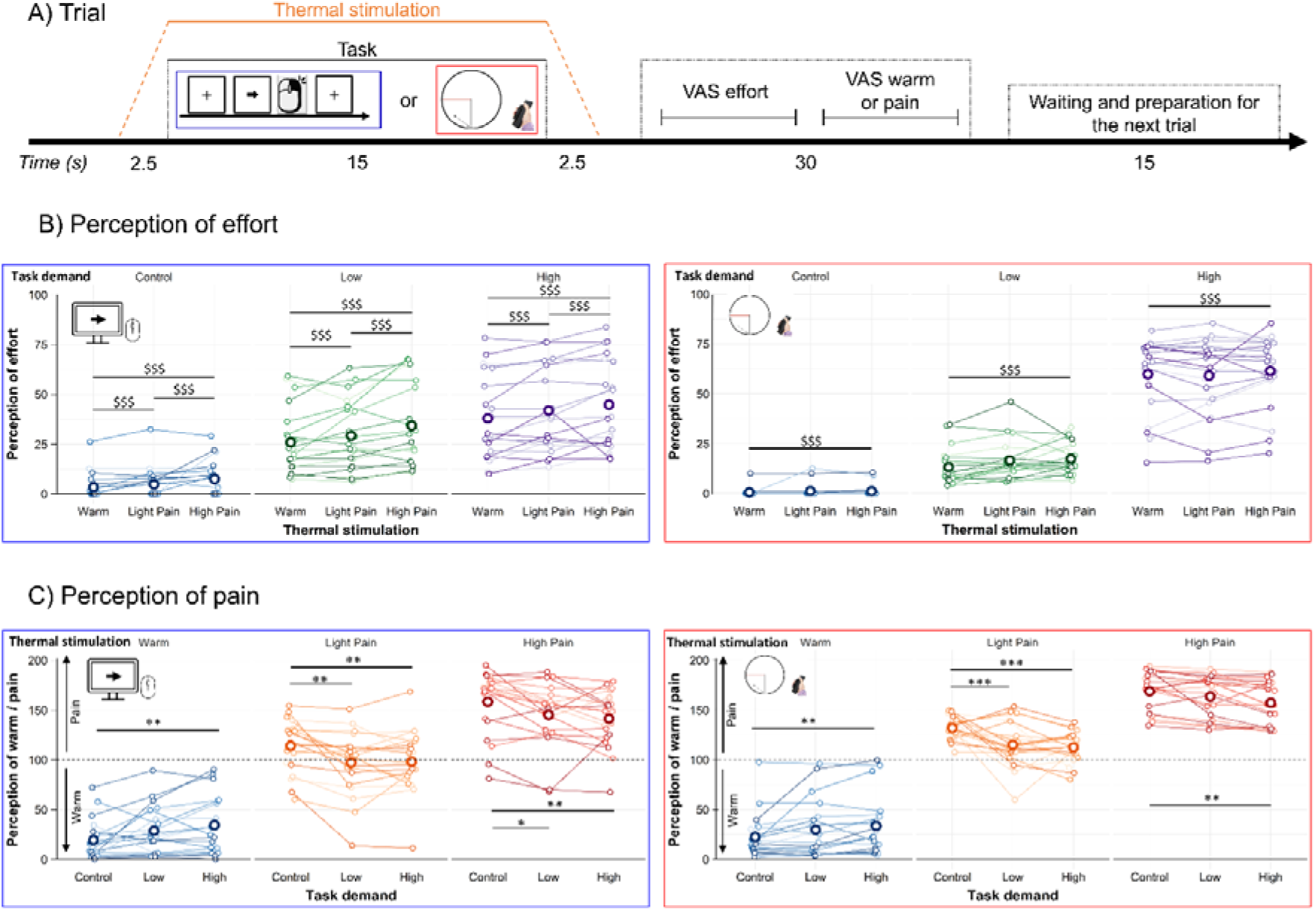
*Panel A* shows the trial sequence: thermal stimulation rises to the target temperature over 2.5 s, remains at plateau for 15 s, and returns to baseline over 2.5 s. During the plateau, participants perform a cognitive or motor task. Immediately afterward, they rate the effort invested with a visual analog scale, report whether the stimulation felt warm or painful, and then rate its intensity with a visual analog scale. *Panels B and C* depict effort and pain perceptions, respectively, as a function of thermal level (panel B) or task demand (panel C), separately for each task demand/thermal stimulation and experiment type (cognitive, blue; motor, red). In both panels, bold circles indicate means, and shaded lines represent individual participants. $ denotes post-hoc tests for main effects and * for interactions: One, two and three symbols for *p*<.05, *p*<.01, *p*<.001, respectively.

## Statistical analyses

### Statistical tests

The statistical analyses were conducted on R (version 4.4.1) using multilevel modeling, with task demand and pain as fixed effects, and participants as a cluster variable. The post-hoc comparisons were performed using the estimated marginal means, with the Holm-Bonferroni correction. Computational analyses were performed using the maximum likelihood estimate, and the optimization function was either Nelder-Mead or BFGS, depending on the convergence of the models. The computational analyses were conducted in two stages. The first stage involved creating simple mathematical models and progressively adding predictors to increase the complexity of the models. We also varied the models by using either task demand or the logarithm of task demand to predict the perception of effort. Additionally, we tested models incorporating either thermal stimulation levels (objective) or perceived pain (subjective) as predictors of effort perception. In the second stage, we tried to model the theoretical framework of motivational intensity theory. For both types of analyses, the Bayesian Information Criterion (BIC) was used to determine which model best fit the data.

## RESULTS

### Performance and effort perception (Figure 1B)

*During the cognitive tasks*, in the presence of pain, participants reported a higher perceived effort [*F*(2, 654) = 26.78, *p* < .001] to maintain cognitive performance [reaction time, *F*(2, 436) = 0.82, *p* = .443; accuracy, *F*(2, 436) = 0.95, *p* = .387]. The perception of effort increased between warm and low pain condition [*t*(654) = 3.27, *p* = .001, β = 2.87], between warm and high pain condition [*t*(654) = 7.31, *p* < .001, β = 6.41], and between low and high pain condition [*t*(654) = 4.04, *p* < .001, β = 3.54].

*During the motor tasks*, in the presence of pain participants reported a higher perceived effort [*F*(2, 664) = 4.72, *p* = .009] to maintain motor performance quantified with the force coefficient of variation [*F*(2, 429.32) < 0.001, *p* > .999]. The perception of effort increased between warm and high pain condition [*t*(664) = 3.07, *p* = .007, β = 2.07], but did not increase between warm and light pain condition [*t*(664) = 1.49, *p* = .227, β = 1.00], neither between low and high pain condition [*t*(664) = 1.59, *p* = .227, β = 1.07].

### Pain perception (Figure 1C)

In both experiments, pain decreased in response to higher task demands [Cognitive: *F*(2, 648) = 5.68, *p* = .004; Motor: *F*(2, 645.01) = 5.98, *p* = .003]. An exploratory analysis revealed an interaction between task demand and thermal stimulation on perception of pain: cognitive experiment *F*(4, 654) = 9.52, *p* < .001; motor experiment *F*(4, 645.01) = 11.76, *p* < .001. *In the cognitive experiment*, in the warm condition, the perception of warmth increased between the control and high task demand condition [*t*(654) = 3.40, *p* = .004, β = 14.95], but not between control and low task demand condition [*t*(654) = 2.10, *p* = .145, β = 9.23], neither between low and high task demand condition [*t*(654) = 1.30, *p* = .580, β = 5.73]. In the light pain condition, pain perception decreased between control and low task demand condition [*t*(654) = -3.81, *p* = .001, β = -16.74], and between the control and high task demand condition [*t*(654) = -3.61, *p* = .002, β = -15.88]. However, pain perception did not change between low and high task demand condition [*t*(654) = 0.20, *p* = .845, β = 0.86]. In the high pain condition, the perception of pain decreased between control and low task demand condition [*t*(654) = -2.99, *p* = .015, β = -13.13], and between control and high task demand condition [*t*(654) = -3.90, *p* = .001, β = -17.15]. However, the pain perception did not change between low and high task demand condition [*t*(654) = -0.92, *p* = .721, β = -4.03]. *In the motor experiment*, in the warm condition, the perception of warmth increased between the control and the high task demand condition [*t*(645) = 3.24, *p* = .008, β = 11.18], but not between control and low task demand condition [*t*(645) = 2.07, *p* = .197, β = 7.18], neither between low and high task demand [*t*(645) = 1.15, *p* = .502, β = 4.00]. In the light pain condition, the pain perception decreased between control and low task demand condition [*t*(645) = -4.67, *p* < .001, β = -16.18], and between the control and high task demand condition [*t*(645) = -5.68, *p* < .001, β = -19.65]. However, the pain perception did not change between low and high task demand condition [*t*(645) = -1.00, *p* = .502, β = -3.47]. In the high pain condition, the perception of pain decreased between control and high task demand condition [*t*(645) = -3.37, *p* = .006, β = -11.66]. However, the pain perception did not change between control and low task demand condition [*t*(645) = -1.54, *p* = .369, β = -5.33], neither between low and high task demand condition [*t*(645) = 1.83, *p* = .269, β = 6.33].

To summary, warmth perception increased with task demand and pain decreased with task demand.

### Modelling of perceived effort (Figure 2)

Computational analyses used maximum likelihood estimation, and model comparisons relied on the Bayesian Information Criterion (BIC). Because Motivational intensity theory predicts a linear relationship between task demand and effort (Richter et al., 2016), whereas classic psychophysical laws such as Weber-Fechner suggest a logarithmic scaling of subjective experience (Dehaene, 2003), we compared, for each candidate model, linear and logarithmic formulations of task demand. We also compared objective thermal stimulation levels with subjective pain ratings, as proposed in our hypothesis. All models predicted trial-by-trial fluctuations in perceived effort. The analyses indicated that the logarithm of task demand better predicted effort perception in 67% of models, while the remaining 32% showed equivalent prediction. Perceived pain consistently (100%) outperformed thermal stimulation as a predictor of effort perception (Figure 2). In the cognitive experiment, the model that fits the best with the data is a formalization of the motivational intensity theory:

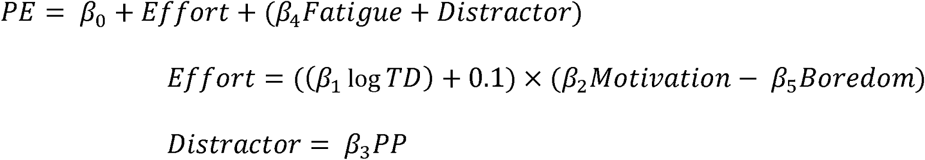

**Figure 2.**
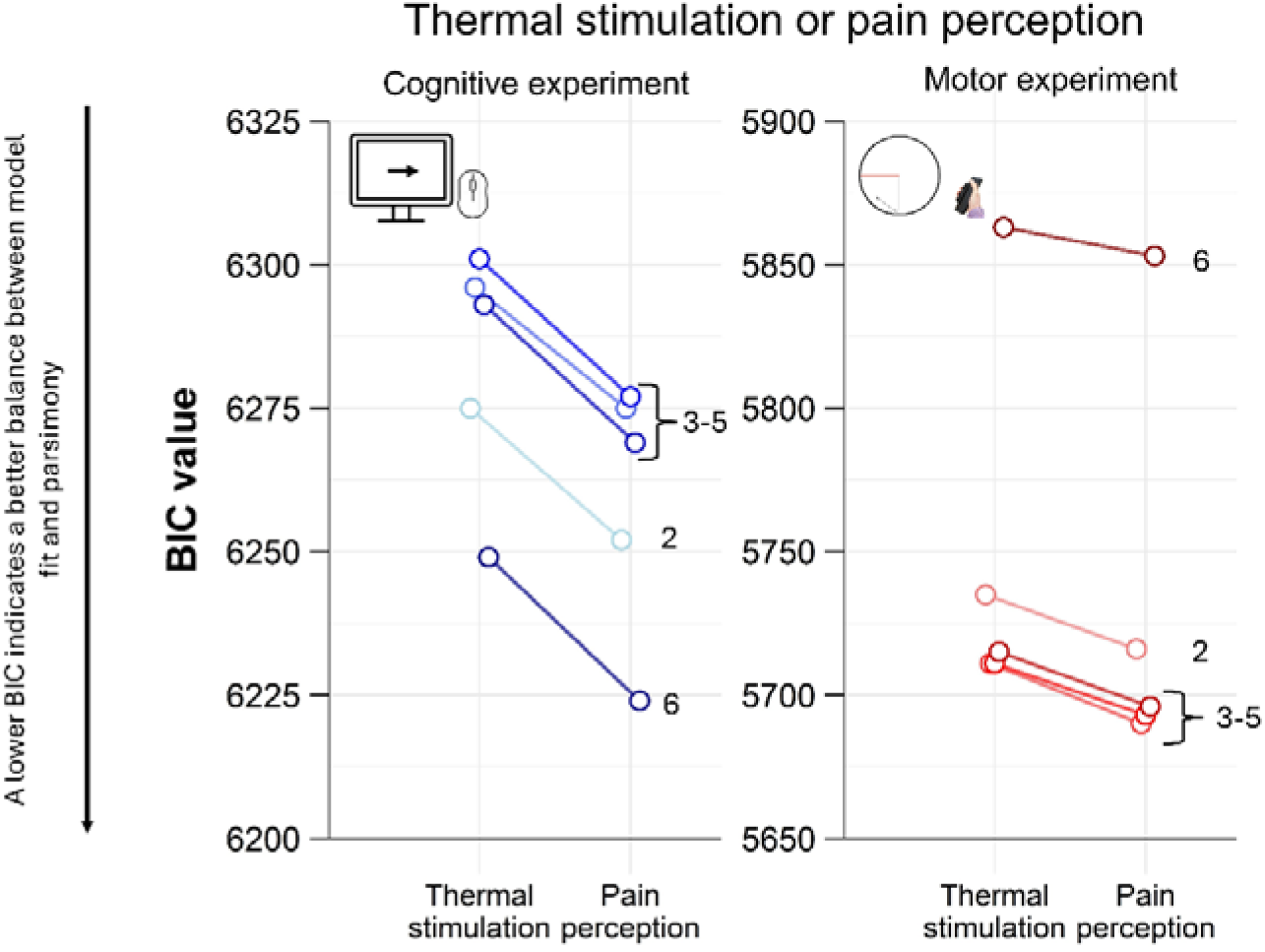
The figure displays Bayesian Information Criterion (BIC) values for the compared models. The figure contrasts models using perceived pain or thermal stimulation to predict effort perception in the cognitive (blue) and motor (red) experiments, indicating which predictor best accounts for the data. Only logarithmic models are displayed here, linear models show the same pattern. Numbers identify the models detailed in the online supplementary information.

Where PE is the perception of effort, TD is the task demand and PP is the perception of pain.

In the motor experiment, the model that fits the best with the data includes the performance:

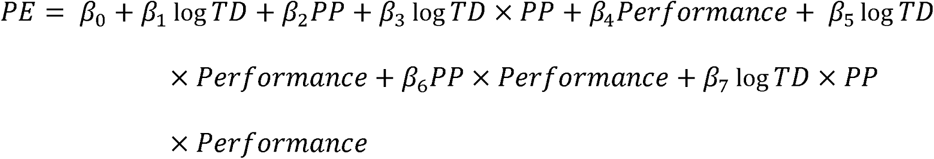

Model specifications and BIC values are provided in online supplementary information.

## DISCUSSION

This study examined how pain interacts with task performance and perceived effort across cognitive and motor domains. We tested whether pain passively degrades performance by diverting resources, or whether performance can be actively preserved through compensatory mechanisms at the cost of increased effort. Across two experiments, our results support the latter account.

Despite pain, performance remained stable across task demands in both cognitive and motor tasks. This preservation was accompanied by increased perceived effort, indicating that participants actively mobilized additional resources to counteract nociceptive interference. These findings argue against a purely passive resource-diversion view and instead support a compensatory account in which effort links the relationship between pain and performance under submaximal demands. The convergence of results across cognitive and motor tasks further suggests a domain-general mechanism. In parallel, in both experiments, pain perception decreased as task demand increased, during light and high pain. This task-induced hypoalgesia indicates that nociceptive processing is actively inhibited during goal-directed action. Critically, the co-occurrence of increased perceived effort and reduced pain supports the interpretation that maintaining performance relies on active cognitive control rather than simple attentional distraction. Effort thus indexes the subjective cost of modulating pain-related interference.

Regarding the increase in perceived warm with increasing task demand observed in both experiments, this effect appears to reflect a *central tendency effect* driven by perceptual uncertainty (Xiang et al., 2021). Indeed, when a warm non-painful stimulus is delivered while participants are performing an effortful task, their attention remains focused on the task rather than being drawn to the thermal stimulation. This interpretation is also consistent with evidence showing that cognitive load, which increases with task demand, enhances the *central tendency effect* (Xiang et al., 2021). Consequently, when participants are subsequently asked to evaluate the stimulation, their ratings tend to shift toward the midpoint of the scale.

Computational analyses provide a formal account of the observed behavioral effects. Trial-by-trial fluctuations in perceived effort were better predicted by subjective pain ratings than by objective thermal stimulation, indicating that effort tracks experienced interference rather than physical nociceptive input. This supports the view that effort reflects the subjective cost of internal regulatory processes, including active pain modulation. Perceived effort scaled logarithmically, rather than linearly, with task demand, consistent with psychophysical principles (Dehaene, 2003) and inverted U-shaped account of effort investment (Silvestrini et al., 2023). This observation places our submaximal tasks on the ascending part of the curve prior to motivational or capacity-limited disengagement, as predicted by the motivational intensity theory (Brehm & Self, 1989; Gendolla & Richter, 2010; Richter et al., 2016). Future studies should test demands beyond these limits to characterize disengagement.

Together, these findings support a unified framework in which effort mediates the relationship between pain and performance. Pain does not inevitably impair task performance but impact depends on the capacity to mobilize compensatory resources. The increased perception of effort reflects the subjective cost of this regulation, while task-induced hypoalgesia reflects its efficacy. By combining behavioral and computational approaches across domains, this study provides converging evidence for effort as a central variable integrating task demand and pain during goal-directed behavior.

## Acknowledgments

We thank Chloé Kos for her assistance with data collection, Todd Vogel for his support with the E-Prime software, and Sean Devine for his help with computational analyses.

TM was supported by the postdoctoral scholarship of the Centre de Recherche de l’Institut Universitaire de Gériatrie de Montréal (CRIUGM), and the postdoctoral research scholarship from the Fonds de Recherche du Québec – Nature et Technologie. IM was supported by Bourse d’études du Réseau québécois de bio-imagerie and Bourse de Mérite aux Cycles Supérieurs de l’Université de Montréal. MM was supported by two Bourses de Recherche du Premier Cycle (BRPC) by the Conseil de Recherches en Sciences Naturelles et en Génie du Canada (CRSNG). MR was supported by Canada Research Chair (tier 2) on Brain Imaging of Chronic and Experimental Pain. BP is supported by the Natural Sciences and Engineering Research Council of Canada – Discovery Grant and the Chercheur Boursier Junior 1 from the Fonds de recherche du Québec – Santé. This project was part of an international collaboration between BP and SB supported by the XIe Commission mixte permanente Québec–Wallonie-Bruxelles.

## Author Contributions

TM, IM, SB, MR, PR and BP designed the study. TM, IM and MM collected the data. TM performed the statistical analyses. TM and BP created the figures. TM and IM wrote the first draft of the manuscript. All authors revised and approved the final version of the manuscript.

## AI Statement

Generative artificial intelligence tools were used during the preparation of this manuscript to assist with the formulation and translation of selected text passages and to support the development of R scripts used in the computational analyses. AI-generated suggestions were used solely as an aid and were not incorporated without critical evaluation. All text, code, analyses, and interpretations were reviewed, validated, and approved by the authors. The authors retain full responsibility for the integrity, accuracy, and originality of the work.

