## Supplementary materials for "Maintaining performance under pain is effortful: experimental and computational evidence"

1 École de kinésiologie et des sciences de l’activité physique (EKSAP), Faculté de médecine, Université de Montréal, Montreal, Canada, QC H3T 1J4

2 Centre de recherche de l’Institut universitaire de gériatrie de Montréal (CRIUGM), Montreal, Canada, QC H3W 1W5

3 Laboratory of Applied Biology, Research Unit in Applied Neurophysiology (LABNeuro), Université Libre de Bruxelles (ULB). Bruxelles, Belgium, 1070

4 Department of Anesthesia, Faculty of Medicine and Health Sciences, McGill University, Montreal, Canada, QC H4A 3J1

5 Alan Edwards Centre for Research on Pain, McGill University, Montreal, Canada, QC H3A 2B4

6 Département de stomatologie, Faculté de médecine dentaire, Université de Montréal, Montreal, Canada, QC H3T 1J4

7 Centre interdisciplinaire de recherche sur le cerveau et l’apprentissage (CIRCA), Montreal, Canada, QC H3T 1P1

* Thomas Mangin

The preregistrations for both experiments can be found respectively at the following links: [cognitive experiment: osf.io/wf8es; motor experiment: osf.io/5hgwk].

Study design

**Participants.** Twenty-one young adults participated in each study. One participant has been excluded from each experiment. During the cognitive experiment, one participant had difficulties staying awake, and closed his eyes between each trial. During the motor experiment, one participant reported not understanding the instructions. Both participants finished the experiment but were not included in the analyses. Final samples were composed of 20 participants for each experiment, including 10 females. Every participant identified their gender according to their sex attributed at birth. The mean (and SD) age for the 20 participants in cognitive and motor experiments is 21.9 (1.6) and 22.2 (1.8) respectively. The weight of the participants is 66.1kg (12.6kg) and 65.4kg (9.9kg) respectively, for a height of 171 cm (11 cm) and 169 cm (8 cm).

**Protocol overview.** Both the motor and cognitive study were composed of two laboratory visits spaced by 24h to one week. Both visits took place at the same time of the day (± 1h) to control the influence of the circadian cycle.

During visit 1 (~2h), participants were asked general questions to control for confounding factors (e.g., substances use such as cigarettes, caffeine, alcohol and drugs, physical activities practiced and the amount of sleep the night preceding the experiment) and they filled out the Pain Catastrophizing Scale Questionnaire. Successively, they went through a sensory calibration of thermal stimulations, a maximal voluntary contraction (MVC) peak force calculation (only for the motor experiment), a ramp-up of effort with familiarization of the visual analog scale (VAS) related to the perceptions of effort, and a familiarization with the motor or cognitive task consisting of only 6 trials.

In visit 2 (~1.5h) participants performed the motor or cognitive task in the presence of thermal stimulation. At the beginning of the visit, temperatures used for the thermal stimulation were validated and adjusted if necessary. Participants then completed either the cognitive or motor task that consisted of 36 15s trials at three levels of difficulty: no demand (control; looking at a fixation cross, or holding the dynamometer), low or high demand (performing a modified Simon task or matching a target at 5% or 30% of their MVC; see Fig1.A). Simultaneously to the execution of each trial, participants received thermal stimulation at one of three individually calibrated temperatures to produce sensations of warmth (control), light or high pain. After each trial, participants reported their perception of the effort required to complete the task, followed by their perception of warmth or pain.

Task

**Sensory calibration of the thermal stimulations.** All the thermal stimuli were delivered using a 3 x 3 cm contact thermode applied to 4 skin sites on the volar surface of the non-dominant forearm (MEDOC, TSA-II Neurosensory Analyser). Stimuli were delivered between 40°C (slightly warm) and 50°C (max) with the speed of the temperature rise and fall adjusted to last 2 s each and a plateau duration of 15 s.

Participants first completed a tolerance test. In this test, the thermal stimulations at increasingly hot temperatures (from 42 to 49° C, increasing by 0.5° C at each trial) were applied to 4 skin sites on the non-dominant forearm of each participant. Subjects were asked to clearly say stop when they reach a temperature that they consider unbearable. This procedure was repeated two times to be sure about the maximal tolerated temperature.

For the sensory calibration, the method of constant stimuli was implemented to determine each participant’s sensitivity to the thermal stimuli. Thermal stimuli were applied at random temperatures (never above the participant’s tolerance) to 4 skin sites on the non-dominant forearm. For each participant, a stimulus-response function was calculated to determine 3 temperatures successively used during each trial of the motor or cognitive task. The 3 temperatures were selected to produce a sensation of warm (no pain; ~30/200 VAS), low pain (~130/200 VAS), and strong pain (~170/200 VAS) on a 0-200 VAS.

**MVC.** The MVC peak torque was collected with a handgrip dynamometer (TSD121B-MRI, BIOPAC) connected to a MP150 system (BIOPAC).

The MVC was recorded with the participant seated on a chair with the elbow at 90° and a wrist brace to maintain the hand aligned with the forearm. Each participant was asked to maintain the handgrip dynamometer in his/her dominant hand and to squeeze the dynamometer as strong as possible to produce a maximal force without losing the arm position. Participants completed 3 MVCs of 3s each with a one-minute rest in between each contraction. The highest peak of the three signal force curves obtained in Acknowledge, represented the subjects’ MVC. The MVC procedure was realized during both the 1^st^ and 2^nd^ visit. For the 1^st^ visit, the MVC was used to set the template of the familiarization with the VAS for the effort perception. For the 2^nd^ visit, particularly of the motor study, the MVC was used to set the three contractions intensities of the motor task: 0% MVC (rest, control condition), 5% MVC and 30% MVC.

**Effort familiarization.** For the cognitive experiment, the ramp-up of effort is made for participants to experiment different level of effort associated with different cognitive demands. Participants performed a series of a modified Simon task with increased difficulty on each trial by either reducing time allowed to answer or adding executive functions in the task (left-right Simon task, left-right with time reduced, left-right Simon task plus around 50% switching condition, left-right Simon task plus around 50% switching condition and reduced time, Random Simon task, Random Simon task with reduced time). To maximize task engagement, each time the participant makes an error or does not respond in time, a sound « bip » will alert the participant. When the most difficult task is reached, the difficulty will decrease until the easiest task. Each trial is interspaced by up to 35s. At the end of each trial, participants will be asked to rate their perception of effort to perform or trying to perform the task.

For the motor experiment, participants were familiarized with the use of the handgrip dynamometer and the visual analog scale (VAS) for rating their perceived effort. They executed a force-matching contraction with the dominant hand at the following intensities in the following order 5%, 15%, 30%, 60%, 80%, 60%, 30%, 15%, 5% of the individual MVC, and rated the perceived effort after each contraction on the VAS (0-100). Each contraction last for 10 s and was interspaced by up to 35s.

**Motor task.** The motor task was a visuomotor force-matching task consisting of executing voluntary submaximal isometric handgrip contractions to match a force feedback line using a handgrip dynamometer (TSD121B-MRI, BIOPAC) connected to a MP150 system (BIOPAC).

The isometric handgrip contractions involved 15s isometric prehensions with the dominant hand at the following intensities: 12 x low force (5% MVC), 12 x high force (30% MVC) and 12 x control (holding the handgrip without producing force). The order of these intensities was randomized. Visual feedback of the force produced during the contraction was provided on the screen, and participants had to match a force target line. To control for the anticipation of the force contraction, the target always remained at the same position on the screen, but more or less force was required to match it, depending on the trials condition. The onset and offset of the muscle contraction was indicated by an auditory signal. For the control condition (i.e., not producing any force), participants were asked to simply hold the handgrip dynamometer, hence there were no auditory signals.

**Cognitive task.** Participants performed 15s of cognitive tasks, answering by clicking on a mouse with their dominant hand at the following cognitive demands: 12 × low demand (left-right Simon task), 12 × high demand (Random Simon task) and 12 × control (fixing a fixation cross). Task completion was performed concomitantly to thermal stimulation applied to the non-used arm. The order of these tasks was randomized. After the task, participants rated their perception of effort to accomplish or trying to accomplish the task, and then their perception of warm or pain induced by the thermal stimulation. Data of these 3 tests was saved on E-Prime 3.0 (version 3.0.3.80).

A trial of the left-right Simon task (low demand) consists of a series of 10 arrows display in the middle of the screen. First a fixation cross appears in the middle of the screen for 250ms. Then an arrow pointing either on the right or left side appears until the participant answers or for a maximal of 1000ms, if the participant does not answer. This is followed by a fixation cross in the middle of the screen for the remaining of the 1000ms of the stimulus presentation plus 250ms. The participants are instructed to click as fast and accurate as possible to indicate the direction of the arrow. If the arrow is pointing toward the right, they have to click on the right button of the mouse, and if the arrow is pointing toward the left, they have to click on the left button. Each trial last 15s.

A trial of the random Simon task (high demand) follows the same time course than the left-right task. However, around 50% (fully randomized) of the first fixation cross is encompass by a circle. This clue indicates to the participants to answer by clicking to the opposite direction of the arrow (if the arrow point to the left, participants must click on the right button). Furthermore, the arrows are not displayed in the middle of the screen but at either 15% or 85% of the screen on the X axis.

**Trials.** During both the motor and cognitive study, participants executed two blocks of 18 trials while receiving concomitant thermal stimulations.

Each trial consisted of 20s thermal stimulation during which participants performed the motor or cognitive task. The motor and cognitive task always started 2.5s after the beginning of the thermal stimulation, meaning at its plateau.

The thermal stimulation and motor or cognitive task were followed by a 40s recovery period used to record perceptions of effort and pain as well as a recovery period to limit the development of fatigue. During the 40s recovery period, participants first rated the intensity of the perceived effort to perform the task on a validated VAS (0-100). Second, they had to indicate if the thermal stimulation was painful or warm but not painful, by choosing between two buttons labelled “pain” and “warm” presented on the screen. Depending on the selected button (pain or warm), participants rated the intensity of the thermal stimulation using a VAS (0-100).

The experimenter moved the thermal stimulator to a different arm spot each trial (4 different arm spots of the non-dominant forearm) to prevent a possible sensitization of the stimulated area. The order of the warm vs low pain vs high pain conditions, as well as the control vs 5% vs 30% MVC conditions were pseudorandomized across trials.

Statistical analyses

**Sensitivity analysis.** For both experiments, the primary rational for our sample size was linked to the financial resources available for our project (Lakens, 2022).

A preliminary pilot study with 7 participants allowed us to determine that the effect size for the interaction on the perception of effort in physical task between Effort (5 vs 50% of the MVC) and Thermal stimulation (41.5, 44, 46.5 and 49°C) was equal to *ƞ²_p_* = .568. However, we think that this effect size is overestimated, especially for perception of effort in mental task. We think that a *ƞ²_p_* around .300 is more realistic.

A sensitivity analyses for a repeated measure ANOVA with “*option as in SPSS*”, alpha risk = .05 and a sample size of 20 and 6 measure indicated that we have 80% of chance to observe an effect size of *f* = 0.378 or higher (which is equal to *ƞ²_p_* of 0.125). Because our estimated effect size is higher, we have a high probability of observing the expected behavioral results.

**Models**

Acronym:

TD = Task Demand

PE = Perception of Effort

PP = Perception of Pain

*Models are the same for both experiments*

To assess motivation, fatigue, and boredom, three self-reported measures were collected: prior to the start of the task, at the midpoint (after 18 trials), and immediately following the final trial. To model the evolution of these states at the level of individual trials, values were linearly interpolated between each pair of successive measurement points. Specifically, the value assigned to trial 1 corresponded to the pre-task rating, while that of trial 18 matched the mid-task rating; intermediate trials were assigned values based on linear progression between these two endpoints. For instance, if a participant reported a fatigue level of 67 before the task and 85 at mid-task, the value increased by (85–67)/17 per trial until trial 18. The same procedure was applied between trials 19 and 36, interpolating between the mid-task and post-task ratings.

Model 1: This is the simplest model that includes only task demand.

Model 1: PE = β0 + β1*TD

Model 1 log: PE = β0 + β1*log(TD)

Model 2: This model adds stimulation (temperature or perception of pain), that allows to compare both on prediction of perception of effort. There is also the interaction between task demand and stimulation.

Model 2 Temperature: PE = β0 + β1* TD + β2*Temperature + β3*TD*Temperature

Model 2 PP: PE = β0 + β1*TD + β2*PP + β3*TD*PP

Model 2 Temperature log: PE = β0 + β1*log(TD) + β2*Temperature + β3*log(TD)*Temperature

Model 2 PP log: PE = β0 + β1*log(TD) + β2*PP + β3*log(TD)*PP

Model 3: This model extends the previous specification by adding *Performance* and its interactions with Task Demand and Temperature as predictors. Performance is conceptually linked to perceived effort. It is generally accepted that performance tends to improve when individuals increase their voluntary effort, particularly in controlled laboratory contexts (2).
In the cognitive experiment, performance is quantified using the inverse efficiency score (IES), which combines response speed and accuracy (3).
In the motor experiment, performance corresponds to the coefficient of variation of force, normalized by the participant’s maximal voluntary contraction (MVC).

Model 3 Temperature: PE = β0 + β1*TD + β2*Temperature + β3*TD*Temperature + β4*Performance + β5*TD*Performance + β6*Performance*Temperature + β7*TD*Performance*Temperature

Model 3 PP: PE = β0 + β1*TD+ β2*PP + β3*TD*PP + β4*Performance + β5*TD*Performance + β6*Performance*PP + β7*TD*Performance*PP

Model 3 Temperature log: PE = β0 + β1*log(TD) + β2*Temperature + β3*log(TD)*Temperature + β4*Performance + β5*log(TD)*Performance + β6*Performance*Temperature + β7*log(TD)*Performance*Temperature

Model 3 PP log: PE = β0 + β1*log(TD) + β2*PP + β3*log(TD)*PP + β4*Performance + β5*log(TD)*Performance + β6*Performance*PP + β7*log(TD)*Performance*PP

Model 4: This model extends the previous specification by adding *Motivation* as an additional predictor of perceived effort, allowing us to account for potential variance in PE attributable to motivational state.

Model 4 Temperature: PE = β0 + β1*Task demand + β2*Temperature + β3*TD*Temperature + β4*Performance + β5*TD*Performance + β6*Performance*Temperature + β7*TD*Performance*Temperature + β8*Motivation

Model 4 PP: PE = β0 + β1*TD+ β2*PP + β3*TD*PP + β4*Performance + β5*TD*Performance + β6*Performance*PP + β7*TD*Performance*PP + β8*Motivation

Model 4 Temperature log: PE = β0 + β1*log(TD) + β2*Temperature + β3*log(TD)*Temperature + β4*Performance + β5*log(TD)*Performance + β6*Performance*Temperature + β7*log(TD)*Performance*Temperature + β8*Motivation

Model 4 PP log: PE = β0 + β1*log(TD) + β2*PP + β3*log(TD)*PP + β4*Performance + β5*log(TD)*Performance + β6*Performance*PP + β7*log(TD)*Performance*PP + β8*Motivation

Model 5: This model extends the model 3 specification by adding *Fatigue* as an additional predictor of perceived effort, allowing us to account for potential variance in PE attributable to fatigue state. Fatigue increases the perceived difficulty of a task, then participant must increase their effort to perform it (4).

Model 5 Temperature: PE = β0 + β1*TD+ β2*Temperature + β3*TD*Temperature + β4*Performance + β5*TD*Performance + β6*Performance*Temperature + β7*TD*Performance*Temperature + β8*Fatigue

Model 5 PP: PE = β0 + β1*TD+ β2*PP + β3*TD*PP + β4*Performance + β5*TD*Performance + β6*Performance*PP + β7*TD*Performance*PP + β8*Fatigue

Model 5 Temperature log: PE = β0 + β1*log(TD) + β2*Temperature + β3*log(TD)*Temperature + β4*Performance + β5*log(TD)*Performance + β6*Performance*Temperature + β7*log(TD)*Performance*Temperature + β8*Fatigue

Model 5 PP log: PE = β0 + β1*log(TD) + β2*PP + β3*log(TD)*PP + β4*Performance + β5*log(TD)*Performance + β6*Performance*PP + β7*log(TD)*Performance*PP + β8*Fatigue

Model 6: This model is a formalization of the motivational intensity theory (4, 5).

Model 6 MIT Temperature: Effort = (β1*TD + 0.1) * (β2*Motivation – β5*Boredom)

Distractor = β3*Temperature

PE = β0 + Effort + (β4*Fatigue + distractor)

Model 6 MIT Temperature log: Effort = (β1*log(TD) + 0.1) * (β2*Motivation – β5*Boredom)

Distractor = β3*Temperature

PE = β0 + Effort + (β4*Fatigue + distractor)

Model 6 MIT Pain: Effort = (β1*TD + 0.1) * (β2*Motivation – β5*Boredom)

Distractor = β3*PP

PE = β0 + Effort + (β4*Fatigue + distractor)

Model 6 MIT Pain log: Effort = (β1*log(TD) + 0.1) * (β2*Motivation – β5*Boredom)

Distractor = β3*PP

PE = β0 + Effort + (β4*Fatigue + distractor)

**The models and the associated BIC**

The red color indicates the winning model for each experiment.

*Cognitive experiment*

Model 1

- Linear version: BIC = 6294
- Logarithm version: BIC = 6276

Model 2

- Linear version and thermal stimulation: BIC = 6292
- Logarithm version and thermal stimulation: BIC = 6275
- Linear version and pain perception: BIC = 6272
- Logarithm version and pain perception: BIC = 6252

Model 3

- Linear version and thermal stimulation: BIC = 6296
- Logarithm version and thermal stimulation: BIC = 6296
- Linear version and pain perception: BIC = 6275
- Logarithm version and pain perception: BIC = 6275

Model 4

- Linear version and thermal stimulation: BIC = 6301
- Logarithm version and thermal stimulation: BIC = 6301
- Linear version and pain perception: BIC = 6277
- Logarithm version and pain perception: BIC = 6277

Model 5

- Linear version and thermal stimulation: BIC = 6293
- Logarithm version and thermal stimulation: BIC = 6293
- Linear version and pain perception: BIC = 6269
- Logarithm version and pain perception: BIC = 6269

Model 6

- Linear version and thermal stimulation: BIC = 6268
- Logarithm version and thermal stimulation: BIC = 6249
- Linear version and pain perception: BIC = 6246
- Logarithm version and pain perception: BIC = 6224

*Motor experiment*

Model 1

- Linear version: BIC = 5732
- Logarithm version: BIC = 5725

Model 2

- Linear version and thermal stimulation: BIC = 5742
- Logarithm version and thermal stimulation: BIC = 5735
- Linear version and pain perception: BIC = 5724
- Logarithm version and pain perception: BIC = 5716

Model 3

- Linear version and thermal stimulation: BIC = 5712
- Logarithm version and thermal stimulation: BIC = 5711
- Linear version and pain perception: BIC = 5692
- Logarithm version and pain perception: BIC = 5690

Model 4

- Linear version and thermal stimulation: BIC = 5713
- Logarithm version and thermal stimulation: BIC = 5711
- Linear version and pain perception: BIC = 5695
- Logarithm version and pain perception: BIC = 5693

Model 5

- Linear version and thermal stimulation: BIC = 5717
- Logarithm version and thermal stimulation: BIC = 5715
- Linear version and pain perception: BIC = 5698
- Logarithm version and pain perception: BIC = 5696

Model 6

- Linear version and thermal stimulation: BIC = 5869
- Logarithm version and thermal stimulation: BIC = 5863
- Linear version and pain perception: BIC = 5859
- Logarithm version and pain perception: BIC = 5853

**Betas values of the winning model for each experiment.**

*Cognitive experiment*

Winning model is model 6 with logarithm of task demand and pain perception.

Effort = (-0.31*log(TD) + 0.1) * (-1.45*Motivation – 0.42*Boredom)

Distractor = 0.07*PP

PE = 2.94 + Effort + (0.13*Fatigue + distractor)

*Motor experiment*

Winning model is model 3 with logarithm of task demand and pain perception.

PE = 0.693 + 184.819*log(TD) + 0.006*PP + 0.126*log(TD)*PP + 0.051*Performance + 7.204*log(TD)*Performance + 0.019*Performance*PP - 0.056*log(TD)*Performance*PP
